# Spatial Transcriptomics-based Identification of Molecular Markers for Nanomedicine Distribution in Tumor Tissue

**DOI:** 10.1101/2022.03.02.482584

**Authors:** Jeong-Bin Park, Jin-Yeong Choi, Hongyoon Choi, Hyung-Jun Im

## Abstract

The intratumoral accumulation of nanomedicine has been considered a passive process, referred to as the enhanced permeability and retention (EPR) effect. Recent studies have suggested that the tumor uptake of nanomedicines follows an energy-dependent pathway rather than being a passive process. Herein, to explore the factor candidates that are associated with nanomedicine tumor uptake, we developed a molecular marker identification platform by integrating microscopic fluorescence images of a nanomedicine distribution with spatial transcriptomics (ST) information. When this approach is applied to PEGylated liposomes, molecular markers related to hypoxia, glucose metabolism and apoptosis can be identified as being related to the intratumoral distribution of the nanomedicine. We expect that our method can be applied to explain the distribution of a wide range of nanomedicines and that the data obtained from this analysis can enhance the precise utilization of nanomedicines.

## Introduction

Nanomedicines hold great promise to improve disease diagnoses and treatments in those with various illnesses, including cancer ^1^, immunological diseases ^2^, and infectious diseases ^3^. Nanomedicine is developed based on nanotechnology and has several advantages over conventional drug platforms. First, it is capable of loading imaging contrast for enhanced diagnostic imaging. Second, surface modifications and decorations of targeting moieties for enhanced drug delivery are possible. Finally, it is relatively simple to load various types of treatment molecules to ensure better therapeutic efficacy ^4^. In 1995, Doxil, a doxorubicin-loaded PEGylated liposome, was approved by the FDA and became the first nanomedicine approved for clinical use ^5^. Currently, there are approximately two dozen FDA-approved nanomedicines, including lipid-based, polymer-based and iron-oxide-based nanoparticles ^4^.

Generally, nanomedicines have a size range of 10-150 nm and demonstrate significantly different pharmacokinetics compared to conventional small-molecule drugs ^6^. Unlike small-molecule drugs, nanomedicines are not freely diffusible into tissues and tend to reside in the vascular space after intravenous administration. In most cases, nanomedicines are removed from circulation through opsonization by serum proteins followed by phagocytosis by the reticuloendothelial system (RES). Various surface modification methods, including PEGylation and the introduction of self-peptides, have been introduced to delay opsonization and thus prolong circulation times. By prolonging the circulation time of nanomedicines, the delivery efficiency can be enhanced. This enhanced delivery of nanomedicines realized by prolonged circulation times is clearly seen in diseased tissue, such as tumors or inflammatory regions but is not prominent in normal tissues ^7, 8^. This phenomenon is called Enhanced Permeability and Retention (EPR), and has become a major theory explaining the improved delivery efficiency of nanomedicines compared to conventional small-molecule drugs. The EPR effect was considered a passive process due to leaky neovascularization and limited lymphatic drainage in diseased tissues compared to normal tissues ^9, 10^.

Recently, the notion that, enhanced tumor accumulation of nanomedicines is a passive process has been challenged. An in vivo imaging study based biodistribution analysis using radiolabeled PEGylated liposomes showed that markers for blood and lymphatic vessel density were not significantly associated with the tumor accumulation levels, in contrast to the prior hypothesis ^11^. Furthermore, the quantified number of endothelial gaps in tumor vasculature is too low to explain the tumor accumulation of the nanoparticles and according to the simulation, 97% of the nanoparticles accumulate in the tumor via an active process ^12^. However, it is very challenging to identify markers that govern active process of the nanomedicine tumor accumulation. So far, only a few potential RNA or protein markers could be analyzed by IHC or reverse transcription polymerase chain reaction (RT-PCR) ^11, 13^. The next-generation sequencing (NGS) technology now can provide an unbiased exploration of the molecular markers. Since tumor uptake of nanoparticles is heterogeneous within the tumor ^14^, it is difficult to find factors that determine nanoparticle uptake by conventional RNA sequencing method that obtains the average value of gene expression in the tissue without spatial information ^15^.

Recent technological advances have established spatial transcriptomics (ST) that can systematically identify the expression levels of all genes throughout the tissue space ^16, 17^. Because ST data inherently possesses spatial information, it can be easily integrated with other types of imaging data and is considered most appropriate for analyzing spatially heterogeneous information within tissues ^17^. Meanwhile, it is possible easily to determine the distribution of the nanomedicine within the tumor by using fluorescently labeled nanomedicine ^18, 19^. Therefore, we hypothesized that molecular factors related to heterogeneous nanomedicine tumor uptake can be identified by the integration of ST data and fluorescent imaging in cancer tissue after the injection of a fluorescently labeled nanomedicine.

## Results and Discussion

### In vivo and ex vivo fluorescence imaging of a mouse xenograft tumor

In this study, a PEGylated liposome was selected as a model nanomedicine, as these liposomes are undoubtedly among the most successful nanomedicine platforms. Also, factors determining the high uptake of PEGylated liposomes remain controversial, as noted in the previous study ^20^. TEM images of PEGylated liposomes showed a uniform and round shape, allowing the identification of a typical lipid bilayer of liposomes. The hydrodynamic sizes of the fluorescent liposomes were 128.05 ± 46.71 nm. The maximum absorbance wavelength and maximum emission wavelength at 550 nm excitation were 550 nm and 563 nm, respectively (**Figure 1A**). Fluorescent liposomes were injected intravenously into 4T1 tumor-bearing mice (n = 3) and in vivo imaging of the mice was obtained using an in vivo fluorescence imaging system (IVIS). We observed that fluorescent liposomes accumulated in the 4T1 tumor in all three mice tested here (**Figure 1B**). Also, ex vivo fluorescence imaging of normal organs and tumors were obtained 24 hours after the injection (**Figure 1C**). Fluorescent signals were observed mainly in the tumors, livers, and spleens. Moreover, the biodistribution pattern of the fluorescent liposomes was similar to the previously reported biodistribution of PEGylated liposomes in tumor-bearing mice ^21^.

**Figure 1.**
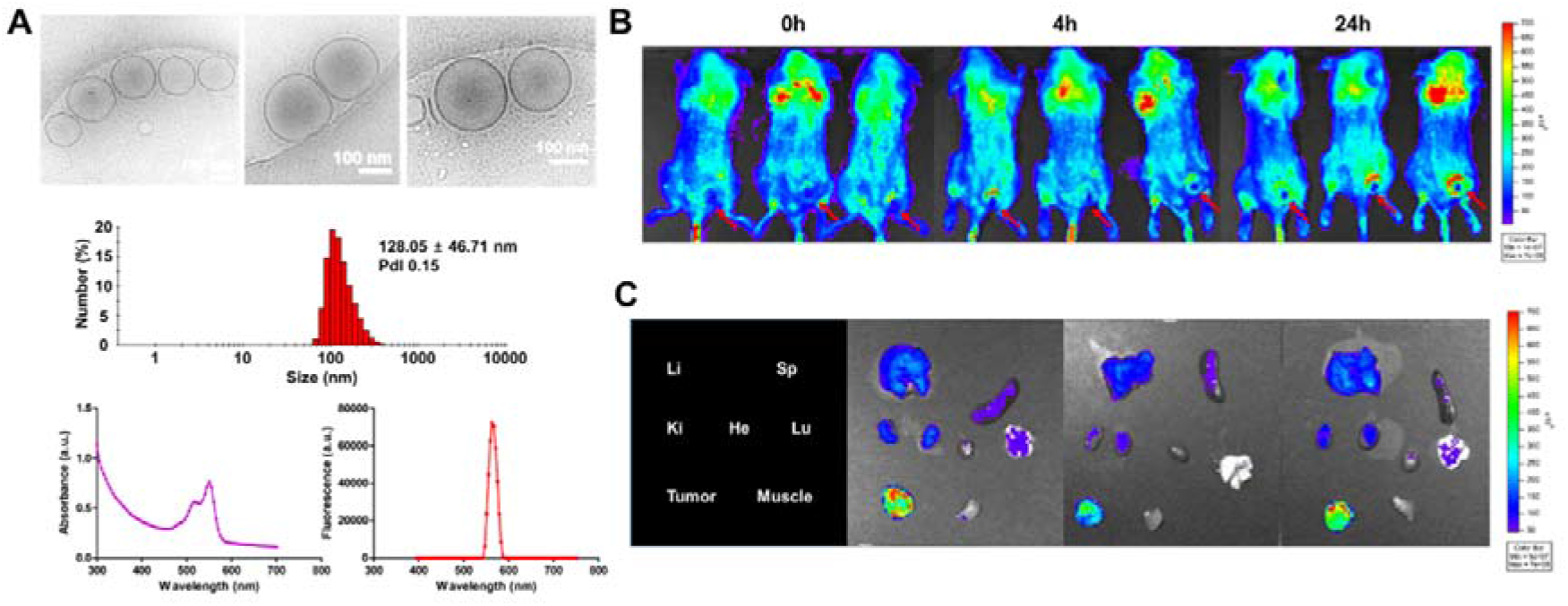
Fluorescence imaging of DiI-loaded liposomes. **(A)** The characterization profile of DiI-loaded liposome using transmission electron microscopy (TEM), dynamic light scattering instrument (DLS) and microplate reader. **(B)** In vivo fluorescence imaging of 0, 4, and 24 hours after intravenous injection in 4T1 breast tumor model. **(C)** Ex vivo fluorescence imaging of main organs (liver, spleen, kidneys, heart, lung, tumor and muscle).

### H&E staining, spatial transcriptomics and fluorescence imaging of the tissue

Among the three excised tumors, the tumor with the highest fluorescent signal was selected for further experiments. We obtained two consecutive sections from the tumor, and the one section was used for H&E staining and ST analysis (**Figure 2A**), and the other was used for fluorescence imaging (fluorescent liposome distribution map) (**Figure 2B**). Spatial mapping of RNA reads indicated a cancer-rich region that showed the highest gene expression, with the necrotic region, the lower and right part of the tissue, showed the lowest (**Supplementary Figure 1**). According to the fluorescence image, the fluorescent signal was prominent in the tumor capsule area, with multiple foci of increased fluorescent signal found in the inner region of the tissue. Next, we obtained a binary map of the fluorescence image, and the map was matched with ST spots for further analyses (**Figure 2B – Figure 2E**). The pattern of the average fluorescence intensity according to the distance was different from the mathematical model for simple passive diffusion (**Figure 2F**). The numerical analysis results of Fick’s law were obtained to predict the passive process of fluorescent liposome distribution using the vascular marker *Pecam1*. The distribution did not concur with the actual florescent liposome distribution and especially could not explain the intra-tumoral uptake of nanoparticles. This was the same with another representative pan-endothelial marker, *Cd34* (**Figure 2B, G-H, Supplementary Figure 2**).

**Figure 2.**
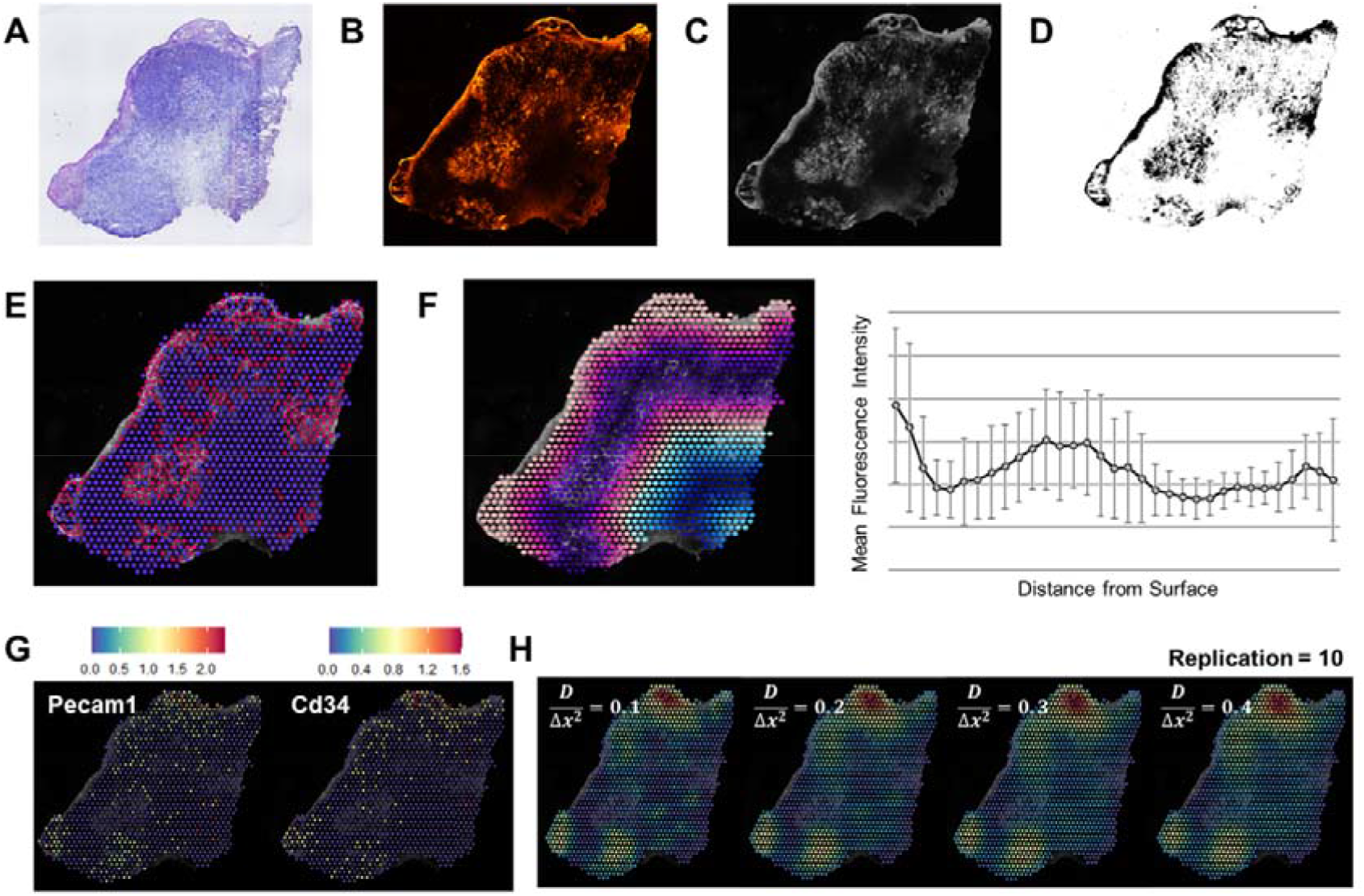
Tumor Sections and Initial Exploration of Data. **(A)** H&E staining image representing the overall histological features. A serial process beginning from original fluorescence image included **(B)** fluorescence image normalization, **(C)** image registration, **(D)** image binarization, and ended with **(E)** acquiring binary map corresponded to the binary image. **(F)** A map colored by the annotated distance and the average fluorescence intensity according to the distance from left-most surface. **(G)** Spatial feature plots of Pecam1 and Cd34 and **(H)** Simulation results of Fick’s diffusion annotated by with different values and numbers of replications.

### Hbb-bs was related to the fluorescence liposome distribution

Firstly, we derived differentially expressed genes (DEGs) related to the high accumulation of fluorescent liposomes in the tumor section by comparing high vs low uptake spots using binary fluorescence imagery. We found that there was one significant gene, *Hbb-bs* (**Figure 3A, Supplementary Table 1**). *Hbb-bs* encodes a beta polypeptide chain found in hemoglobin in red blood cells (RBC) and considered as one of the RBC markers ^22^. Consistently, we found that the expressions of vascular markers (e.g., *Pecam1, Cd34*) were similar to the distribution of *Hbb-bs*. This result implied that the fluorescent liposome distribution is associated with blood circulation (**Figure 3A**).

**Figure 3.**
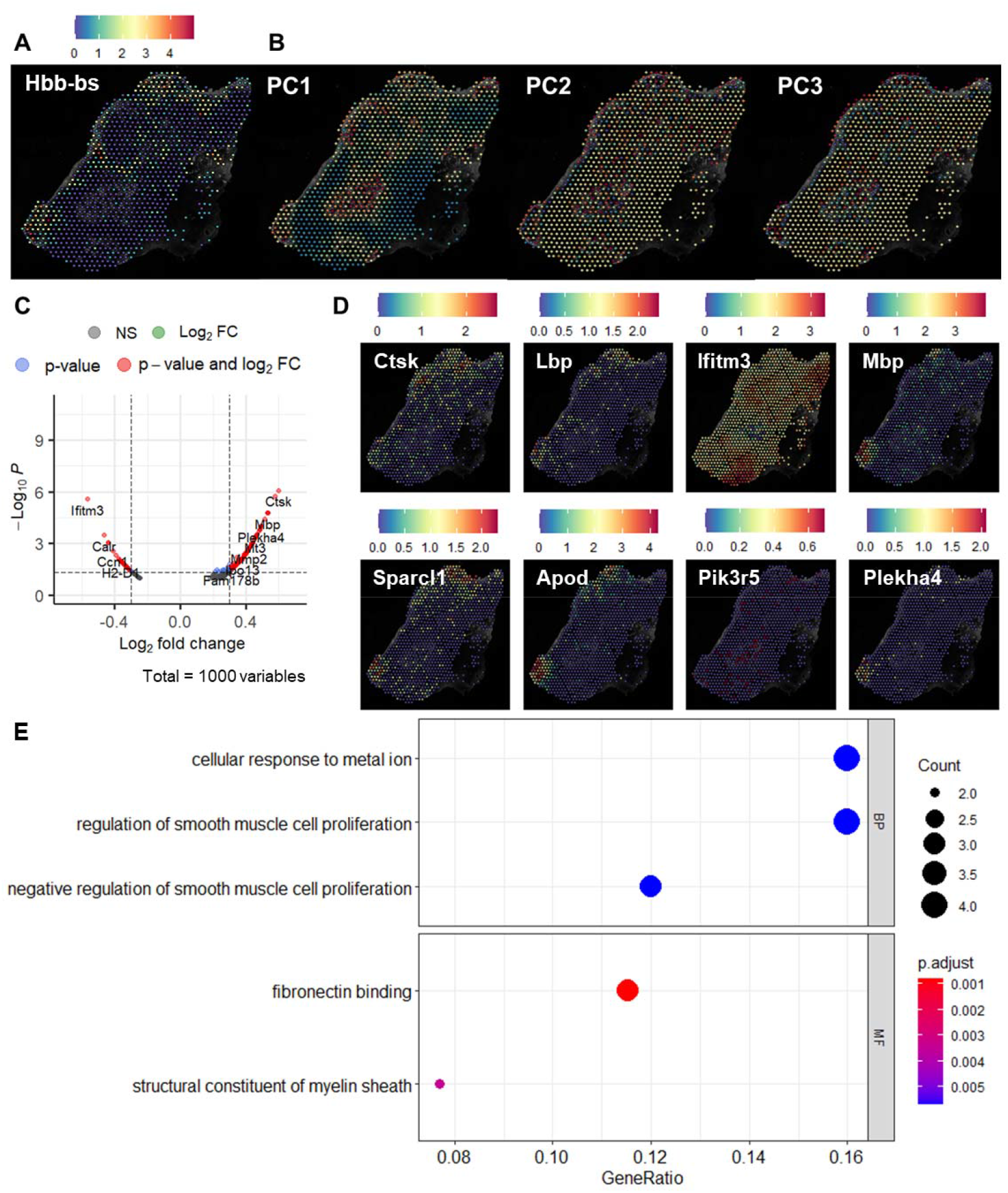
Results of overall fluorescence analysis. **(A)** A spatial feature plot of the only differentially expressed genes (DEG), *Hbb-bs*. **(B)** Image latent features generated by SPADE and PCA algorithm. PC1, PC2, and PC3 referred to principal component 1, and so on. **(C)** Enhanced volcano plot with top 1000 variable genes. **(D)** Spatial feature plots of top 8 SPADE genes with highest fold change (FC). **(E)** GO analysis for PC1 SPADE genes of top 30 up-regulated genes according to biological process (BP), cellular component (CC), and molecular function (MF). Top 3 positive and negative GO terms for each category was considered.

We determined the principal components (PC1, PC2, and PC3) in the fluorescent image of the tumor tissue using spatial gene expression patterns by deep learning of tissue images (SPADE) algorithm ^23^. The most variable latent feature of the fluorescence image (i.e., PC1) showed a pattern similar to that of the distribution of fluorescent liposomes (**Figure 3B**). We obtained PC1-associated genes (SPADE genes) and demonstrated them using enhanced volcano plots (**Figure 3C**). Among the SPADE genes, it is noteworthy that *Ctsk, Lbp, Sparcl1*, and *Apod* were high-ranked and up-regulated genes, which are abundant in the extracellular matrices (ECMs) of the stromal region (**Figure 3D, Supplementary Table 2**). This finding is in line with the previous observation that nanoparticle tumor uptake is associated with capillary wall collagen ^13^. Also, it is well known that *Apod* can be found in the early stage of tumor development among the apolipoproteins, consistent with the present experimental condition ^24^. The SPADE genes were associated with regulation of smooth muscle cell proliferation and fibronectin binding according to the gene ontology analysis (**Figure 3E**). To further investigate the association between the expression of *Hbb-bs* and the nanomedicine distribution, we conducted a correlation analysis of the fluorescent signal intensity and expression level of *Hbb-bs* within high-uptake spots. There was no statistically significant correlation between *Hbb-bs* expression level and fluorescence intensity (r= 0.073, p-value = 0.188).

### Division of uptake pattern clusters and cell type analysis

*HBB-bs* and SPADE genes were found to be related to the fluorescent distribution of nanomedicines in the tissue. However, the expression pattern of the genes did not match the inner uptake clusters of the tumor (**Figure 2B vs. Figure 3A, and 3D**). We speculated that the NP uptake mechanisms on the surface and in the inner area of the tumor could differ, therefore the uptake patterns were analyzed using image feature-based clusters. The regions of fluorescent image were clustered using a CNN-based features based on tiles of the image and K-means clustering. K was set to 4 as the minimum requirement for the division of the peripheral area from the inner area (**Supplementary Figure 3**). Also, we obtained two clusters of high-uptake spots (cluster 1: surface cluster, cluster 2: inner cluster) by matching the regions with the binary map (**Figure 4A**). Through the unsupervised hierarchical clustering of spots in clusters 1 and 2, and outliers were eliminated (**Supplementary Figure 4**)

**Figure 4.**
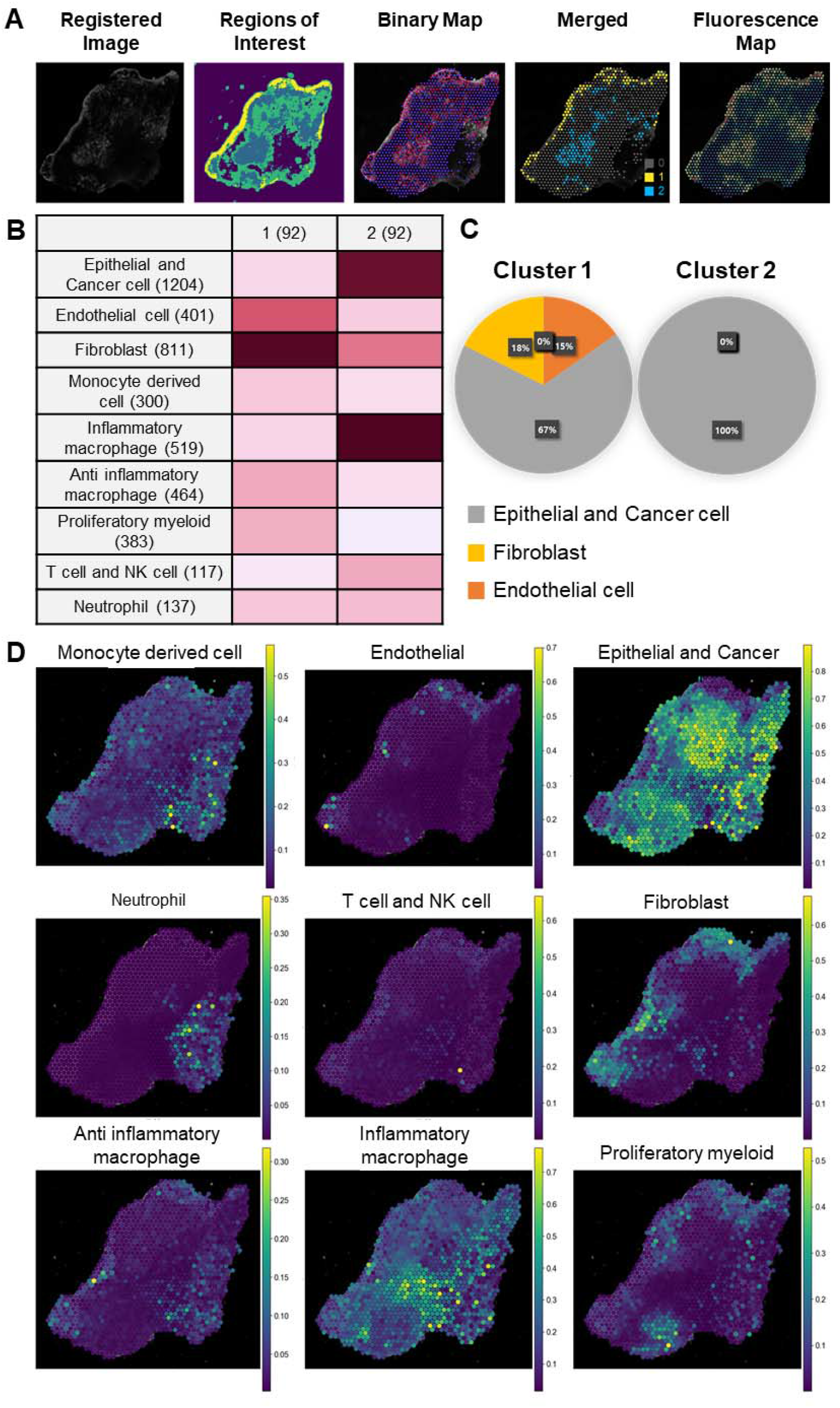
Results of subgroup fluorescence analysis. **(A)** A process determining uptake clusters beginning from registered fluorescence image. **(B)** MIA analysis for cluster 1 and 2. **(C)** Results of cell type population derived from RCTD algorithm. **(D)** Spatial feature plots representing each cell type occurrence acquired from the results of CellDART algorithm.

We then analyzed the cell distribution of the clusters using three different types of cell-type prediction methods, in this case multimodal intersection analysis (MIA) ^25^, robust cell type decomposition (RCTD) ^26^, and single-cell and spatial transcriptomic data (CellDART) ^27^. Using MIA, fibroblasts and endothelial cells were preferentially discovered in cluster 1 while cancer cells were found to be predominant in cluster 2 (**Figure 4B**). In the RCTD analysis, cancer cells were dominant in both clusters and endothelial cells and fibroblasts were the major cell types in cluster 1. We could confirm the relatively dominant distributions of endothelial cells and fibroblasts in cluster 1 compared to cluster 2, an outcome similar to the results of the MIA analysis (**Figure 4C**). Results of CellDART verified the observations from the MIA analysis and RCTD assessment (**Figure 4D**). Cancer cells are dominant in the tumor tissue while endothelial cells and fibroblasts are clearly observed in the surface region of the tumor according to CellDART. Also, the presence of inflammatory macrophages from the MIA assay and the dominance of cancer cells in the RCTD assay could be reconciled by the CellDART results.

### Identification of DEGs and uptake-associated genes in cluster 1 and 2

We conducted a DEG analysis of clusters 1 and 2 by comparing cluster 0 vs 1 and 0 vs 2. Volcano plots of DEGs showed different genetic profiles between clusters 1 and 2 (**Figures 5A and B and Supplementary Tables 3-4**). In addition, a dot plot representing the top 20 genes for each cluster verified the uniqueness of clusters 1 and 2 (**Figure 5C**). DEGs of cluster 1 were similar to the result of the previous analysis using the entire tissue slide. For example, we could observe RBC marker such as *Hbb-bs, Hba-a1*, and *Hba-a2* and stromal genes such as *Apod, Aqp1, Col3a1, Gpx3, Apoe*, and *Sparc11* among the top 20 DEGs in cluster 1 (**Supplementary Table 3**)

**Figure 5.**
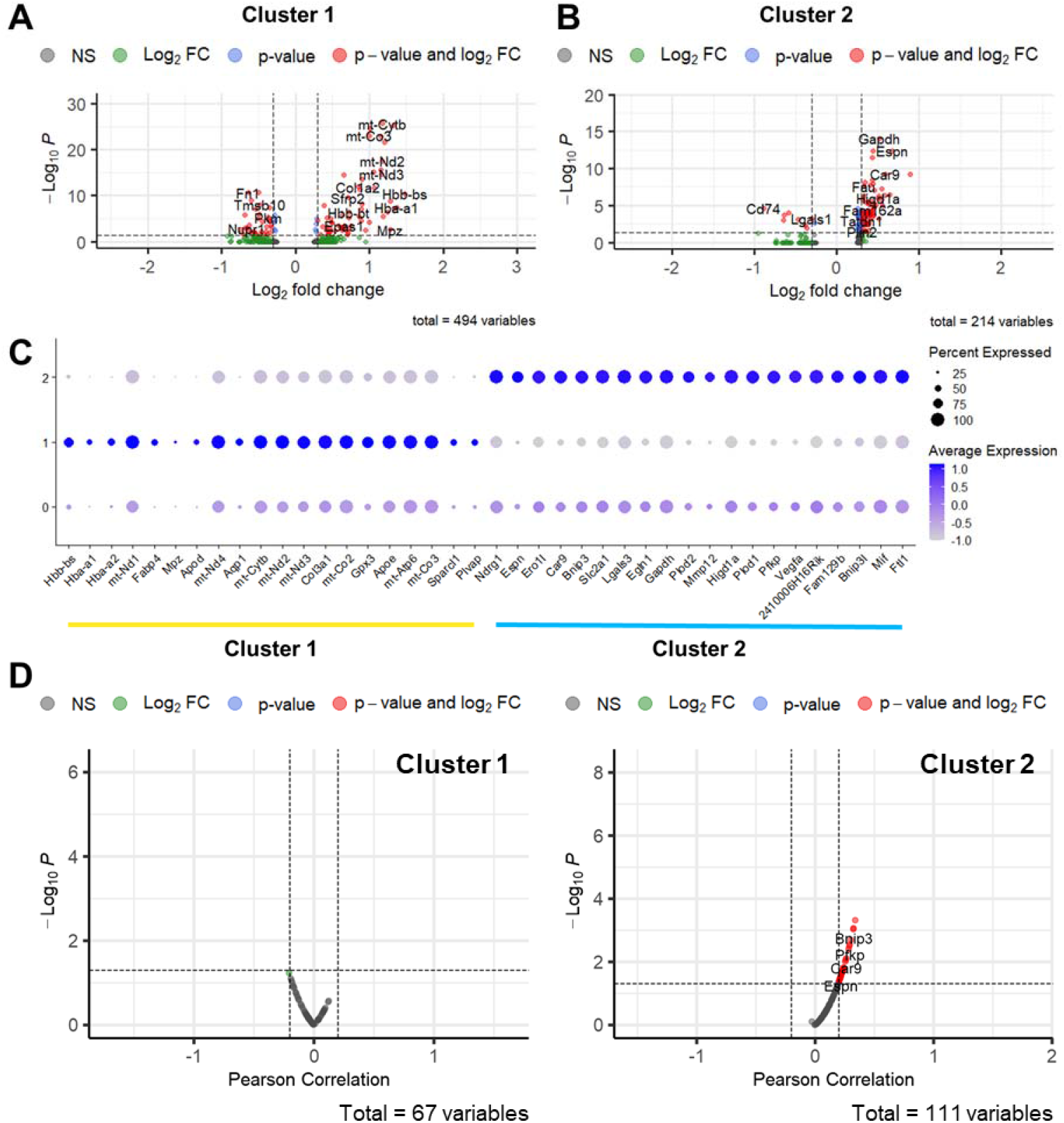
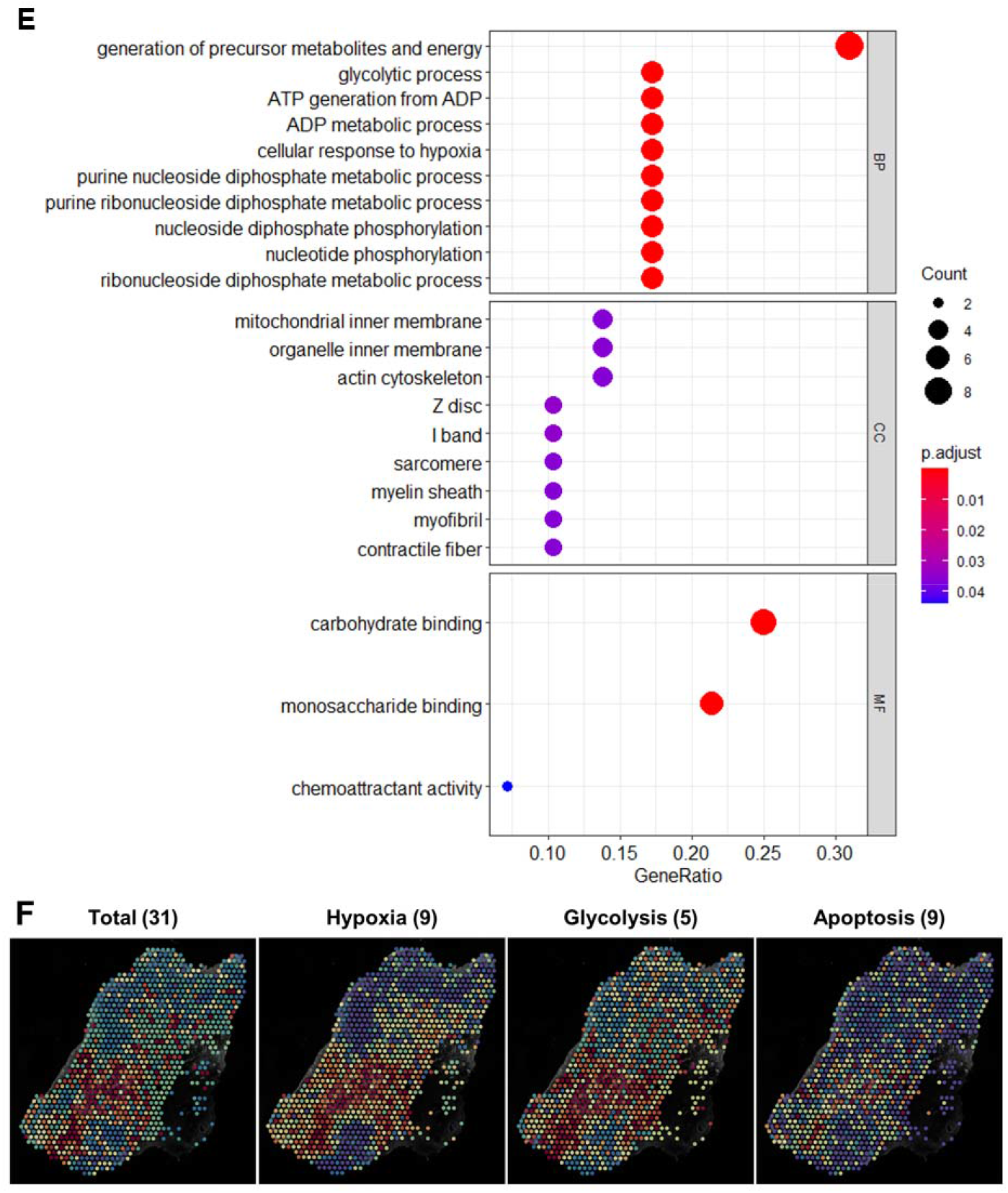
Discovery of uptake-driving genes. Volcano plots of uptake region-specific genes of clusters **(A)** 1 and **(B)** 2. **(C)** A dot plot representing expression of top 20 DEGs of clusters 1 and 2. **(D)** Volcano plots showing the relationship between correlation coefficient and p-value in clusters 1 and 2. **(E)** GO analysis for DEGs with significant correlation of cluster 2 according to biological process (BP), cellular component (CC), and molecular function (MF). **(F)** Spatial feature plots of scores derived from total genes, hypoxic genes, glycolytic genes, and apoptotic genes. The scores were calculated by *AddModuleScore* in Seurat package.

The DEGs in cluster 1 showed no significant correlation with fluorescent signal (**Figure 5D)**. On the other hand, a total of 31 genes that showed upregulated in the cluster 2 and positively correlated with fluorescence intensity in the cluster were identified (**Figure 5D**, **Supplementary Figure 5, Supplementary Table 5**). These genes were associated with three representative physiological functions in gene ontology: hypoxia, glucose metabolism, and apoptosis (**Figure 5E, Supplementary Table 5**). When plotting the scores of hypoxia, glucose metabolism, and apoptosis, the spatial distribution of the scores were all colocalized (**Figure 5F**), and the distribution was similar to the internal distribution of the nanoparticles (**Figure 2B**).

The DEGs from the clusters may simply be region-associated genes because the spots in the clusters are spatially close. However, we speculated that the DEGs with statistically significant correlations with fluorescence intensity within the cluster are associated with nanomedicine uptake process. Several previous studies have reported that a hypoxic condition can enhance NP uptake in cancer cells ^28, 29, 30^. According to our unbiased methods, markers related to hypoxia can be identified to be associated with the PEGylated liposome tumor distribution. We found that the most important glycolysis mediator gene, *Pfkp*, appeared to be an uptake-associated gene in cluster 2, as well (**Supplementary Tables 4 and 5**). It is well known that a hypoxic condition in cancer tissue enhances the glycolysis of cancer cells ^31^. Also, a previous study showed that a significant correlation existed between the degree of hypoxia and glucose metabolism as assessed by in vivo positron emission tomography (PET) in patients’ tumors ^32^. Because a lack of energy generation is prevalent in cancer cells due to the low efficiency of hypoxic metabolism, starvation-induced apoptosis can be triggered ^33^. Also, molecular markers related to lipid metabolism were found to be uptake-driving genes. Hypoxia can reprogram a number of genes related to energy metabolism. In recent years, a link between hypoxia and lipid metabolism was also revealed. In particular, endocytosis of lipoproteins is enhanced by the upregulation of lipoprotein receptor-related protein (LRP1) ^34^ and very-low-density lipoprotein receptor (VLDLR) ^35^. Thus, we speculated that several hypoxia-induced metabolism-related genes, such as *Ndrg1*, which participates in lipid metabolism, including LDL receptor trafficking, play an important role in the uptake of PEGylated liposomes. In addition, *Plin2*, one of the DEGs in cluster 2 (FC = 0.311197, adjusted p val = 0.034946, cor = 0.177015, p val for cor = 0.073657), may be linked to this speculation, as the gene is involved in the hypoxia-inducible lipid droplet-associated protein with Hif-1α. Finally, we conducted a correlation analysis of the uptake-driving genes to find an association between the genes (**Supplementary Figure 6**). Most of the genes were clustered as sets A1, A2, and A3, which are correspondingly related to ‘hypoxia + glycolysis’, ‘hypoxia + apoptosis’, and ‘apoptosis only’. Also, a heatmap indicated that most genes showed strong connectivity, possibly due to a common biological context, except for the three genes in cluster B.

Heterogeneity in EPR-mediated nanomedicine delivery is considered to be a cause of heterogeneous outcomes in clinical trials of nanomedicines. To improve clinical outcomes, predictive biomarkers for the EPR effect should be established. Therefore, molecular markers to predict the efficiency of EPR are urgently needed to design successful clinical trials. Currently, several methods are suggested as EPR markers, including companion imaging biomarkers using radiolabeled nanoparticles and serum markers related to tumor stroma ^36, 37^. We believe that the molecular markers found by our platform can be used as precise EPR markers after validation in other types of cancer models. Also, the molecular markers found in this study can be utilized to enhance the EPR effect using a gene-drug interaction database. For example, the iLINCS database (http://www.ilincs.org/ilincs/) provides a list of small-molecule drugs that enhance or inhibit the molecular process when the molecular markers are provided. The enhancer identified by the database can be used to enhance the EPR effect of the nanomedicine. Taken together, the molecular markers found by the platform here can optimize the precise utilization of nanomedicines by predicting the EPR effect and finding EPR enhancers for the nanomedicines.

In this study, we tested only one example nanomedicine in one tumor model, PEGylated liposome and the 4T1 tumor model, respectively. The tissue distribution of a nanomedicine can be affected by materials, surface modifications and possibly by internal drug loads. Therefore, the result from this study cannot be generalized to other types of nanomedicine. However, if there is a target nanomedicine and target cancer tissue, this new method can then be used to optimize the component and surface chemistry of the nanomedicine to obtain a better tissue distribution pattern. In addition, a three-dimensional or temporal approach should be investigated later for more accurate conclusions.

## Conclusion

Herein, we developed a molecular marker identification platform for nanomedicine distributions by integrating ST data and fluorescence nanomedicine distribution imagery. The molecular markers related to hypoxia, glucose metabolism, lipid metabolism and apoptosis are associated with the intratumoral distribution of PEGylated liposome. An interdisciplinary approach including image processing, an AI-algorithm-based gene analysis, biological annotations, and flexible interpretations of complicated mass transfer events were all involved in the development of the marker identification platform. We believe that this platform can be applied to a variety of tissues and other types of drugs, such as peptides, antibodies, and antibody drug conjugates, for the exploration of novel molecular markers related to drug distributions. Moreover, the use of molecular markers can enhance the precise utilization of drug candidates.

## Materials and Methods

### Materials

1,2-Distearoyl-*sn*-glycero-3-phosphocholine (DSPC), cholesterol, 1,1’-dioctadecyl-3,3,3’,3’-tetramethylindocarbocyanine perchlorate were purchased from Sigma-Aldrich, Korea. 1,2-Distearoyl-*sn*-glycero-3-phosphoethanolamine conjugated polyethylene glycol (DSPE-PEG) was purchased from Creative PEGworks. Avanti Mini Extruder was purchased from Avanti Polar Lipids.

### Synthesis and characterization of DiI-loaded liposomes

Liposomes were synthesized by extrusion method with Avanti mini extruder. The liposomes were composed of DSPC, DSPE-PEG, cholesterol, and DiI fluorescent dye (λ_ex_ = 553 nm, λ_em_ = 570 nm). Thin-film lipids were prepared by vaporizing organic solvents and hydrated by distilled water. Hydrated fluorescent liposome layers were extruded using 400 nm and 200 nm pore size membrane filter in order. The hydrodynamic size of DiI-loaded liposomes was about 130 nm in a dynamic light scattering instrument (DLS).

### 4T1 breast cancer model and fluorescence imaging

To prepare a 4T1 allograft tumor model, 4T1 breast cancer cells (10^6^ cells/0.02 mL) were injected subcutaneously into the BALB/c mice at the right thigh region. After 10 days, DiI-loaded liposomes were injected intravenously. In vivo fluorescence imaging was performed at 0, 4, and 24 hours after injection using in vivo imaging system. For the verification of liposome distribution in organs, mice were sacrificed 24 hours after injection. The main organs (heart, lung, kidney, liver, spleen, muscle, and tumor) were collected and observed by in vivo imaging system for fluorescence imaging.

### *Acquisition of spatial transcriptomics (ST) library*, H&E staining image, and fluorescence image

Fresh tumor samples were embedded in the mold with optimal cutting temperature (OCT) compound for cryo-sectioning. ST library was acquired by several steps: cryo-sectioning, fixation, permeabilization, cDNA synthesis, and RNA sequencing. All the methods were carried out in the way that 10x Genomics visium protocol recommended. The fresh tissue samples were embedded in OCT compound (25608-930, VWR, USA). We prepared consecutive tissue slices were used for H&E staining, spatial transcriptomics library, and fluorescence imaging. The slices were acquired by thin blades used in cryotome so that the fluorescence pattern affected by gene expression could be explored thoroughly. The tissue section for ST were placed on Visum slides (both Visium Tissue Optimization Slides, 1000193, 10X Genomics, USA and Visium Spatial Gene Expression Slides, 1000184, 10X Genomics). The fixation was performed under the recommended protocol using chilled methanol. cDNA libraries were prepared and were sequenced on a NovaSeq 6000 System S1200 (Illumina, USA) at a sequencing depth up to 250M read-pairs.

Raw FASTQ files and H&E images were processed by sample with the Space Ranger v1.1.0 software. The process uses STAR v.2.5.1b (Dobin et al., 2013) for genome alignment, against the Cell Ranger (mouse mm10 reference package). The process was performed by ‘spaceranger count’ commend. Notably, to avoid confusion in terms, “pixel” was used only in the fluorescence image and “spot” was used only in the spatial transcriptomics profile. Also, all the following data analysis methods were summarized. (**Table 1**)

**Table 1.** Summary of the analysis procedures.

**Step 3. Acquisition of marker genes**
|  | **<u>DEG analysis</u>** | **<u>SPADE algorithm</u>** |
| --- | --- | --- |
| **<u># of steps</u>** | 3; high-uptake spots configuration → DEG analysis → correlation analysis | 1; SPADE algorithm |
| **<u>Spot value</u>** | The average of fluorescence intensities of spot-size patches | Principal components of features extracted from VGG16 |
| **<u>Correlation</u>** | Pearson correlation coefficient | Empirical Bayes algorithm and linear regression analysis |
| **<u>P-value</u>** | Wilcoxon rank sum test for uptake region-specific genes; Linear regression analysis for uptake-driving genes | Empirical Bayes algorithm and linear regression analysis for SPADE genes |

**Step 4. Acquisition of uptake clusters**
|  | **<u>Combination of VGG16 and K means clustering</u>** |
| --- | --- |
| **<u>Mechanism</u>** | Texture recognition |
| **<u># of steps</u>** | 3; high-uptake spots configuration → VGG16 & K means clustering → acquisition of uptake-clusters |

### Image registration

To fit the shape of the acquired fluorescence image to the spatial transcriptomics spots, the image registration process based on symmetric diffeomorphic registration ^38^ implemented by Python-based open-source DiPY package (**Supplementary Material 1**). The fluorescence image was changed to gray-scale using opencv2 package. For the registration, linear rigid transformation was performed after the matching center of masses of both images. The rigid and affine transformation processes were optimized using mutual information between two gray-scale images. After the linear transformation, nonlinear warping process based on symmetric diffeomorphic registration algorithm was performed using function ‘SymmetricDiffeomorphicRegistration’ with ‘CCMetric’ for optimization. The transformed image was visually evaluated.

### Annotation of distance in spatial transcriptomics spots

We came up with the distance from the surface of the tumor and defined the distance of spots in the left-boundary region as 0. As we raised the distance one by one, we immediately marked the next layer. And then, the map was colored by the annotated distance. Meanwhile, the fluorescence intensity value for each spot was averaged from the corresponding area in the registered fluorescence image. The fluorescence intensity values were averaged according to the distance and a plot showing the relationship between the annotated distance and the average fluorescence intensity was created.

### Mathematical simulation of diffusion

When illustrating dynamics of diffusion, Fick’s law is used typically as follows:

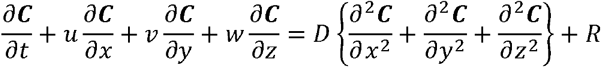

where *C* is concentration vector, (*u, v, w*) is velocity vector, *D* is diffusivity and *R* is source or sink term. We put two assumptions to simulate the result of the formula. At first, we anticipated the number of gaps of blood vessels is proportional to the expression level of *Pecam1* gene. And we thought the tissue sample was close to the median plane of the whole tumor, so we got

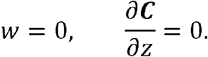

We regarded (*u, v, w*) as null vector according to the literature ^12^. Briefly, when dealing with cylindrical blood vessels perpendicular to the median plane, the maximum fluid velocity in vicinity to the vessels could be drawn with the experimentally estimated value:

*D* (Diameter of blood vessel) = 10 μm
*L* (Spot to spot distance) = 100 μm
*H* (Spot height) = unknown
*S* (Surface area of blood vessels per unit volume of tissue) = 0.0034 μm^2^/μm^3^
*n* (Number of gaps per unit area of blood vessel) = 500 gaps/mm^2^
*V* (Experimentally estimated fluid velocity) = 0.065μm^3^ /s · gap
*V_max_* (Maximum fluid velocity) = unknown
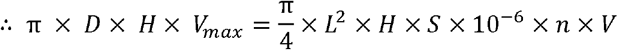

The result of *V_max_* = 27.625 pm/s indicated that the flow rate could not even explain the movement of nanoparticles over tens of millimeters over 24 hours.

Thus, we used a numerical approach to comprehend the effects of the concentration gradient in the absence of the flow rate:

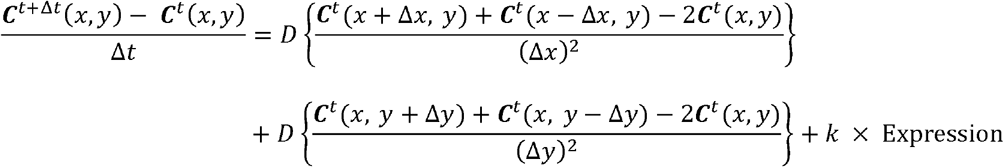

where Δ*x* and Δ*y* were set to be the same. And various simulation results of Fick’s diffusion annotated by *C/k* with different *D*/Δ*x*^2^ values and numbers of replications were explored. (**Supplementary Material 2**)

### Differentially expressed genes based on fluorescence intensity

Image binarization was performed to analyze the fluorescence image pixels combined with the spots with transcriptomic data. Spots with high fluorescence (high-uptake spots) were distinguished from spots with low fluorescence (low-uptake spots) by dichotomy. When making a binary image, only pixels with a brightness greater than 25% of the maximum fluorescence intensity were selected as high pixels. The fluorescence intensity was measured and analyzed by ImageJ (ver 1.8; https://imagej.nih.gov/ij/download.html). Once the binary image was acquired, the binary map was then created in search of the pixel values (i.e., 0 or 1) corresponding to the center of the spots.

To visualize the spots according to gene features, t-distributed stochastic neighbor embedding (t-SNE) was employed using the Seurat package (version 4.0.5) in R. Spots with RNA reads less than 500 were excluded for the following analysis. The perplexity of *RunTSNE* was set to 30. Differentially expressed genes (DEGs) between high-uptake spots and low-uptake spots were explored by Wilcoxon rank sum test on *FindAllMarkers* in Seurat package with *min.pct* and *logfc.threshold* set to 0.25 both. Lastly, DEGs were sorted by fold change (FC). (**Supplementary Material 3**)

### Identification of genes associated with image features

In addition to the overall DEG analysis, another approach, spatial gene expression patterns by deep learning of tissue images (SPADE), was used. (**Supplementary Material 4**) ^23^. A pre-trained VGG16 model extracted 512 features per patch around each spot and principal component analysis (PCA) was performed to reduce the dimensions of the features. We selected top three principal components (PCs) to identify SPADE genes. SPADE genes in each PC were discovered on empirical Bayes algorithm and linear regression analysis. Genes were then sorted by FC.

For visualizing image feature-associated genes, *EnhancedVolcano* function was used with pCutoff of 0.05 and FCcutoff of 0.3. Top 1000 genes with FDRs below 0.05 were selected, and spatial feature plots of top 8 genes with highest FC were represented. For GO analysis, *enrichGO* function was used. GO analysis was performed according to biological process (BP), cellular component (CC), and molecular function (MF) using top 30 up-regulated or down-regulated genes. When specifying biological annotations, g:Profiler (https://biit.cs.ut.ee/gprofiler/gost) was used instead of GO analysis.

### Subgroup fluorescence analysis

We split the fluorescence image into 394 × 384 patches of 5 × 5 patch size and extracted 512 features per each patch by using the VGG16 model. And then patches were classified by K means clustering according to the features with K of 4. (**Supplementary Material 5**) As a result, the fluorescence image was separated into 4 regions of interest (ROIs) according to the texture. Then, the binary map was merged into 2 notable ROIs out of 4. As a result, high-uptake spots were separated into 2 clusters and the other spots were allocated to default 0. In conclusion, 3 clusters were formed totally. Outliers of each uptake cluster were eliminated by using unsupervised hierarchical clustering of spots in cluster 1 or 2. (**Supplementary Material 6**) DEGs representing each uptake clusters were generated by comparison to the default cluster using *FindAllMarkers* in R.

### Analysis of cell types associated with fluorescence distribution

Multimodal intersection analysis (MIA) was performed to comprehend which cell type was relevant to each uptake cluster ^25^. Single-cell RNA sequencing (scRNA-seq) dataset was obtained from the previous research of 4T1 tumor model ^39^. Marker genes in each cell type were determined with adjusted p-value < 1E-05 in Wilcoxon rank-sum test from *FindAllMarkers*. Also, marker genes in each uptake cluster were determined similarly, except for adjusted p-value < 0.01 instead of adjusted p-value < 1×10^−5^. Afterward, each common set of genes in a specific cell type and a specific spatial region was calculated, and enrichment or depletion was determined by comparison with randomized output. Finally, all the common sets of genes were analyzed using Fisher’s exact test to figure out which cell type was significantly characteristic to which spatial region. (**Supplementary Material 7**)

Other cell-type matching algorithms were addressed for verification. Robust cell type decomposition (RCTD) was used to determine the distribution of each cell type through supervised maximum likelihood estimation as a representative alternative method for MIA analysis ^26^. All the parameters were set to the default settings including *doublet_mode* of ‘*doublet*’ in run.RCTD (**Supplementary Material 8**). Another algorithm, CellDART ^27^, which used adversarial domain adaptation classification from single-cell data with pre-labeled cell types was additionally performed to find cell types related to the distribution of fluorescence (**Supplementary Material 9**).

### Genes correlated with fluorescence intensity of subclusters defined by the image

To acquire uptake-driving genes from uptake region-specific genes, the calculation of the Pearson correlation coefficient was performed. We correlated the expression of the gene and the fluorescence intensity within the cluster. As an effect size of the relationship between fluorescence intensity and gene expression, the slope of the regression curve was estimated. Only genes with p value less than 0.05 were sorted according to the correlation coefficient. The resultant uptake-driving genes were explored with gene ontology analysis (**Supplementary Material 10**). The uptake-driving genes were robustly calculated regardless of the criteria for obtaining fluorescence intensity (max, mean+2s.d., mean+1s.d., mean, median, min, etc.), so we decided to calculate it by averaging pixels in spot-size patches without further justification. We used two-step approach, DEG identification at first and correlation analysis later, because unreliable genes easily came up if only correlation analysis was performed.

