## Supplementary Figure 1 to 6 for "Spatial Transcriptomics-based Identification of Molecular Markers for Nanomedicine Distribution in Tumor Tissue"

**Supplementary information**

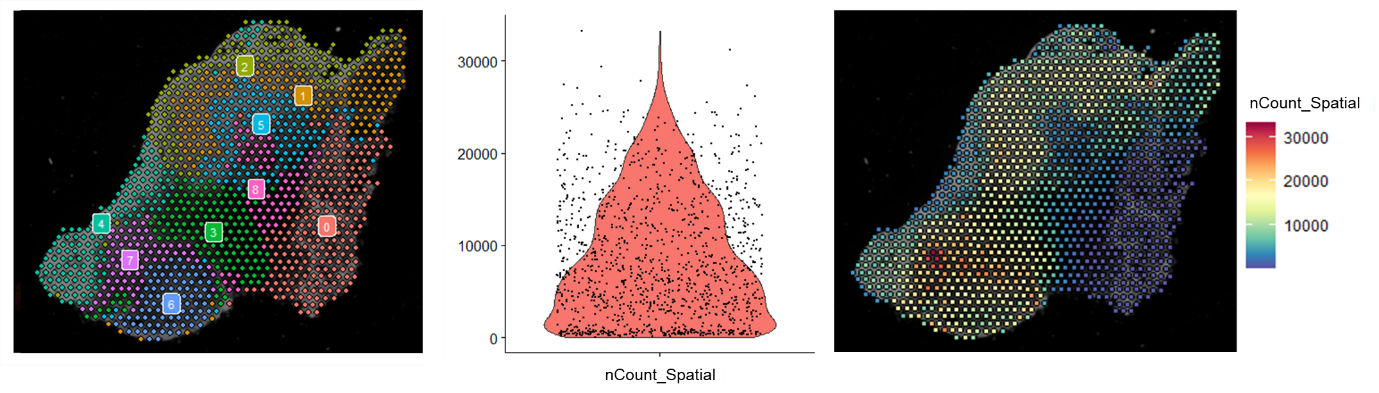

**Supplementary Figure 1. Spatial mapping of RNA reads.**

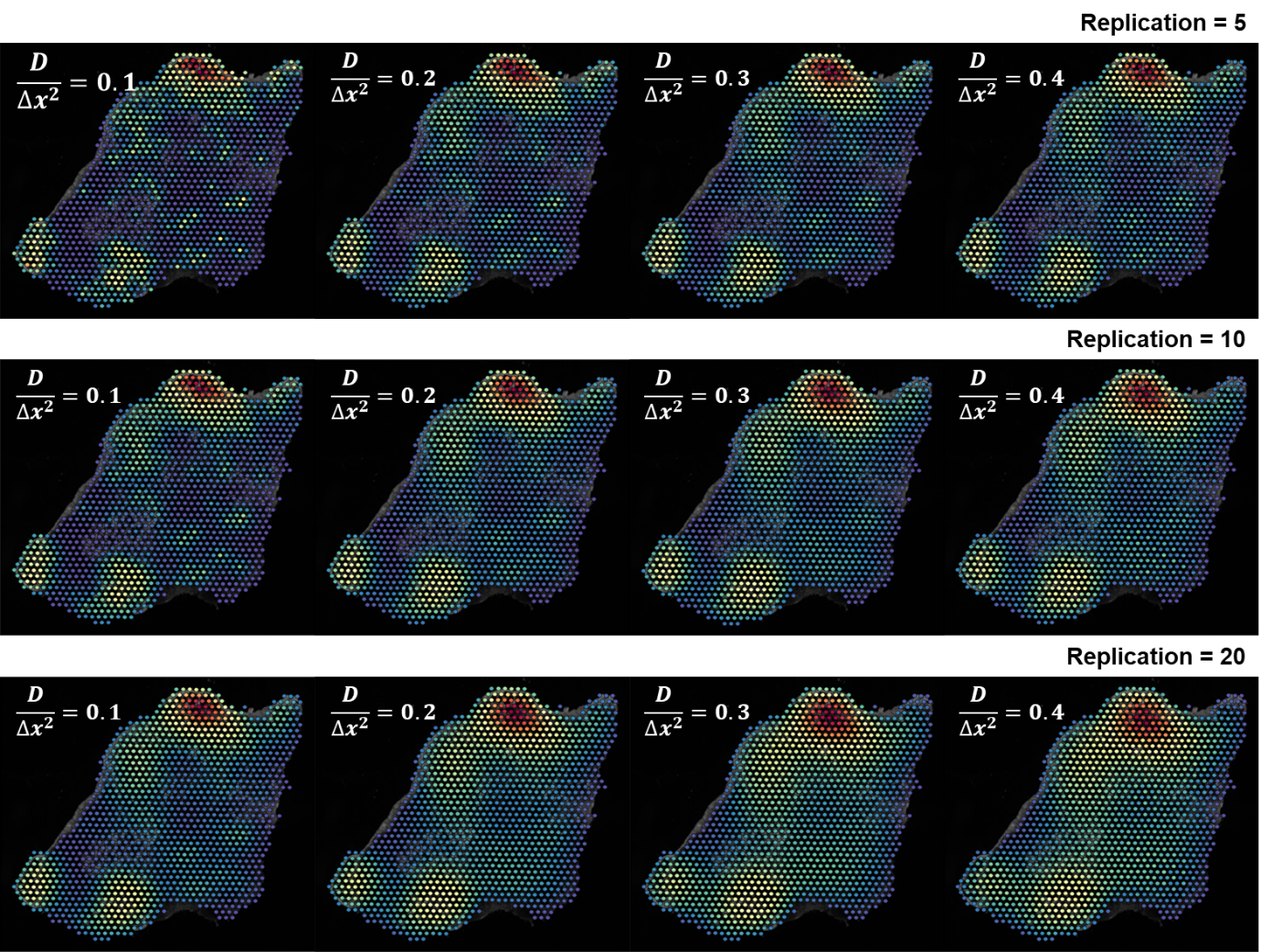

**Supplementary Figure 2. Spatial feature plots of simulation results.** Simulation results of Fick’s diffusion annotated by $C/k$ with different $D/{\Delta x^{2}}$ values and numbers of replications.

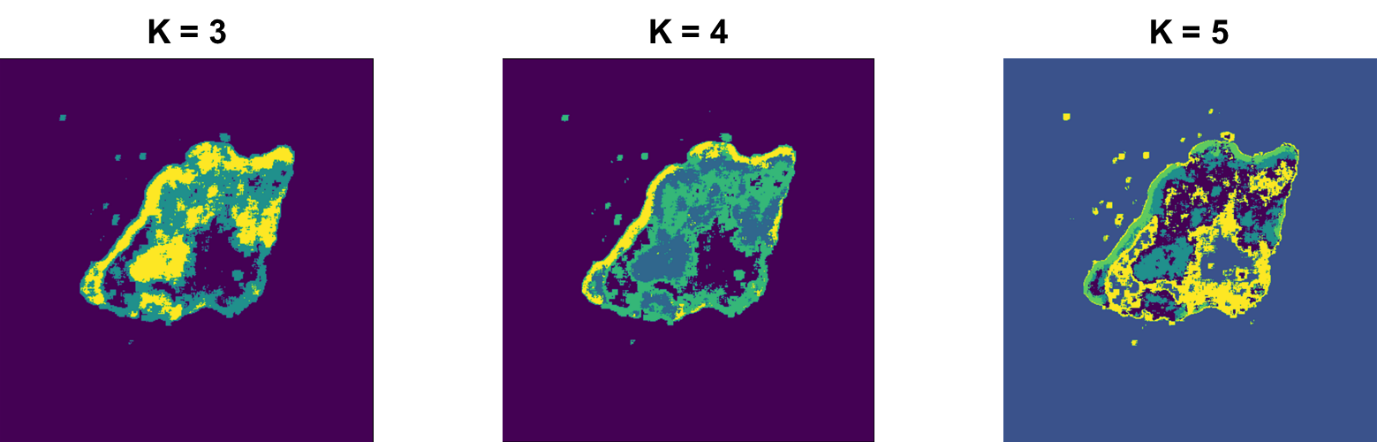

**Supplementary Figure 3. Determination of parameter K in K means clustering.** K was set to 4 as the minimum requirement for the division of the peripheral area from the inner area.

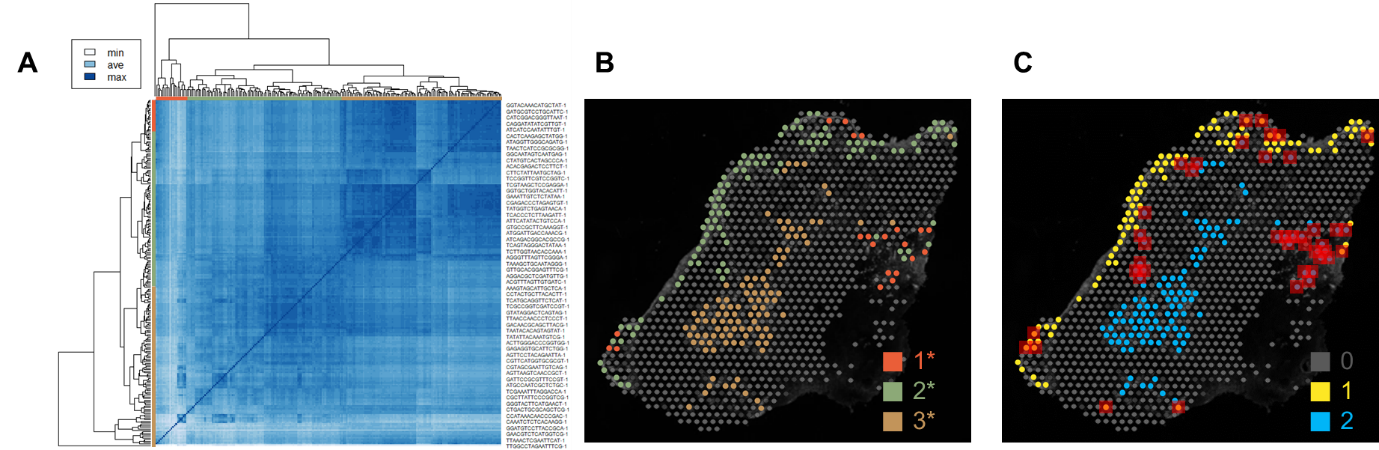

**Supplementary Figure 4. Elimination of outliers in each uptake cluster.** **(A)** A result of unsupervised hierarchical clustering of spots in clusters 1 and 2, and **(B)** spatial feature plots of the spots in clusters 1 and 2 according to the clusters. (C) Identification of outliers in each cluster. Outliers were demarcated by red squares.

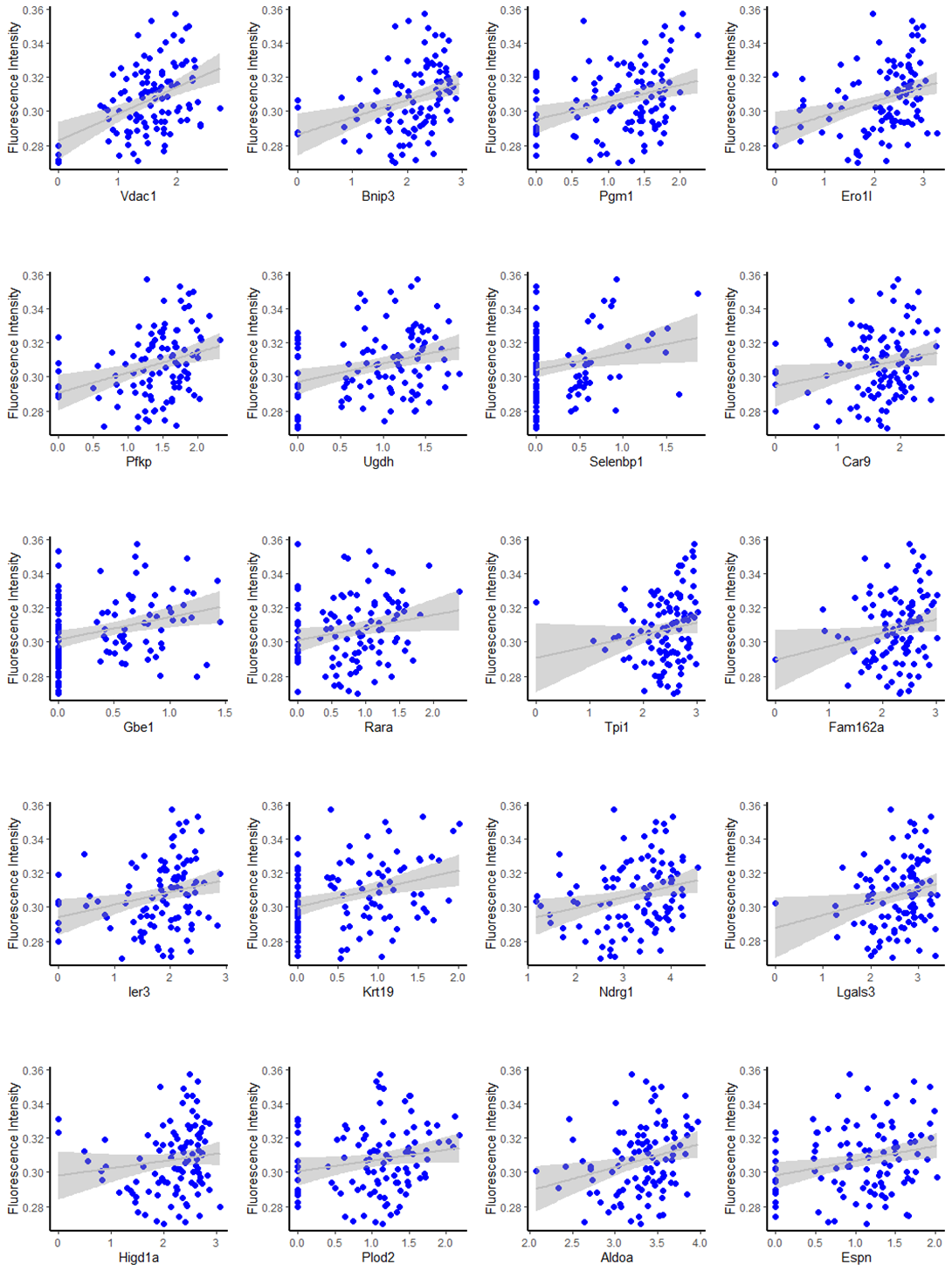

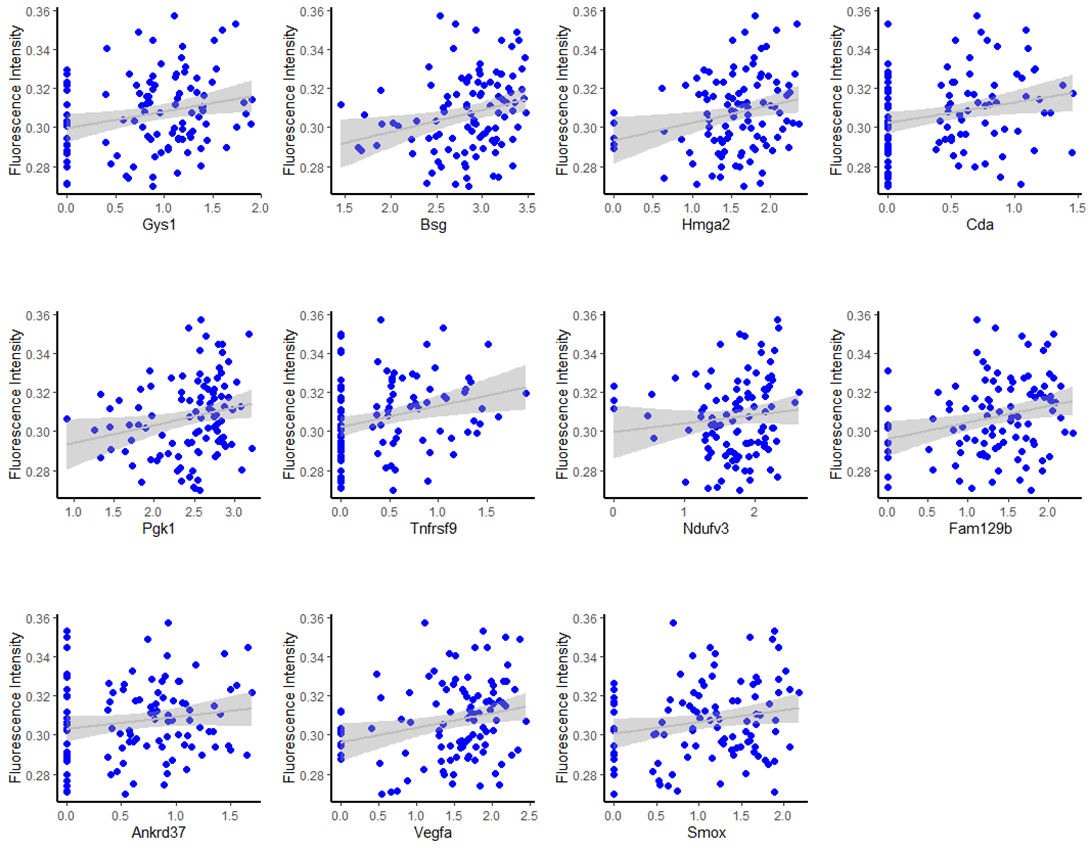

**Supplementary Figure 5. Scatter plots of all uptake-specific genes in cluster 2.**

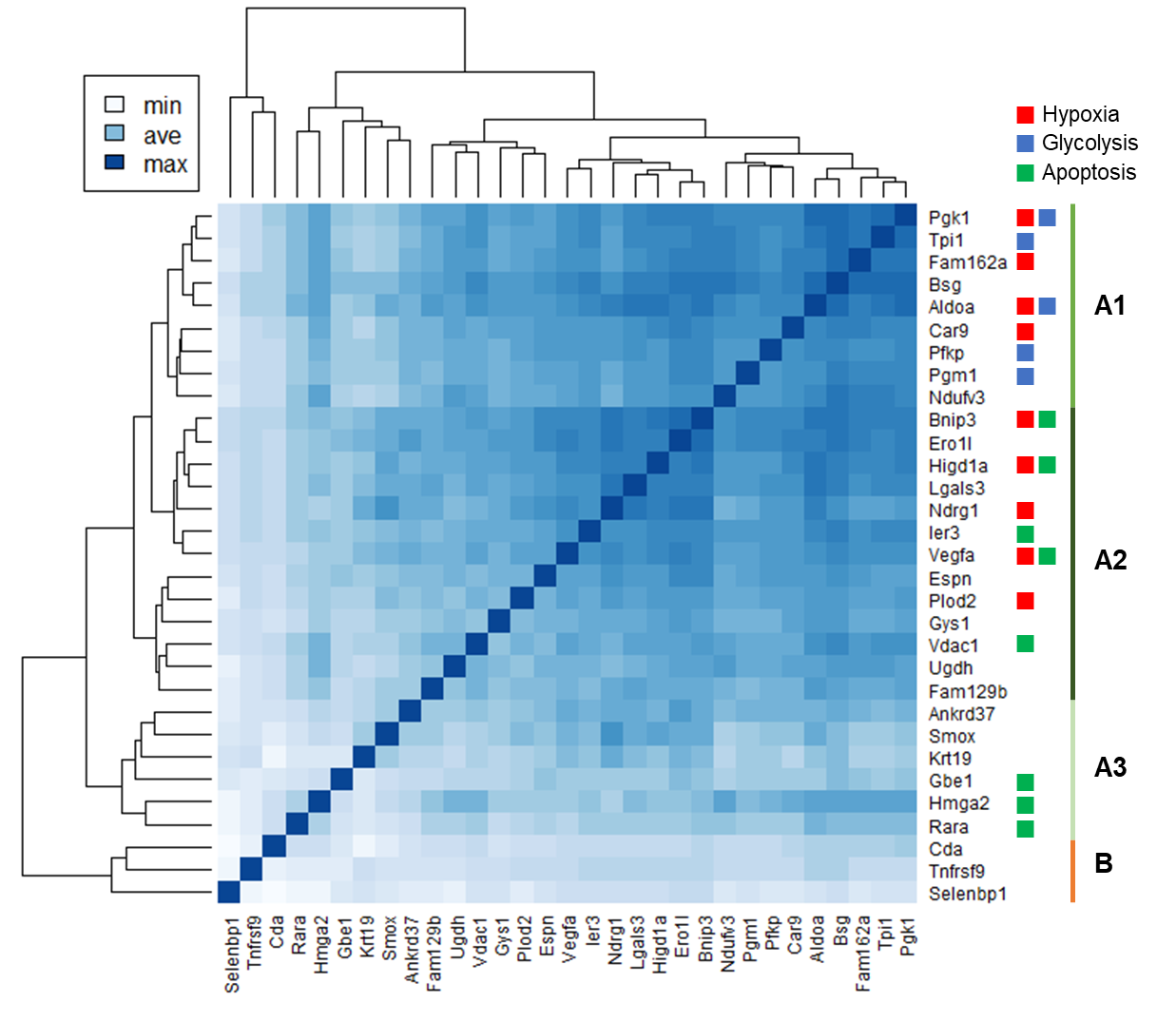

**Supplementary Figure 6. A heatmap derived from the correlation of each pair of DEGs with significant correlation in cluster 2.** The annotations of ‘Hypoxia’, ‘Glycolysis’, and ‘Apoptosis’ were based on the result of g:Profiler about the 31 uptake-specific genes.
