## Supplementary Table 1 to 5 for "Spatial Transcriptomics-based Identification of Molecular Markers for Nanomedicine Distribution in Tumor Tissue"

**Supplementary information**

**Supplementary Table 1. List of top 20 DEGs sorted by FC in overall fluorescence analysis.**

| **Order** | **Gene** | **log2FC** | **p_val_adj** | **Order** | **Gene** | **log2FC** | **p_val_adj** |
| --- | --- | --- | --- | --- | --- | --- | --- |
| 1 | Hbb-bs | 0.806090 | 0.001102 |  |  |  |  |

**Supplementary Table 2. List of top 20 SPADE genes sorted by FC in overall fluorescence analysis.**

| **Order** | **Gene** | **log2FC** | **p_val_adj** | **Order** | **Gene** | **log2FC** | **p_val_adj** |
| --- | --- | --- | --- | --- | --- | --- | --- |
| 1 | Ctsk | 0.597908 | 8.46E-07 | 11 | Fabp4 | 0.483804 | 0.000151 |
| 2 | Lbp | 0.578055 | 1.81E-06 | 12 | Fosb | 0.482819 | 0.000151 |
| 3 | Mbp | 0.534974 | 1.71E-05 | 13 | Aqp1 | 0.478601 | 0.000180 |
| 4 | Sparcl1 | 0.532259 | 1.71E-05 | 14 | Rgs5 | 0.468288 | 0.000309 |
| 5 | Apod | 0.529634 | 1.71E-05 | 15 | Igfbp5 | 0.465283 | 0.000332 |
| 6 | Pik3r5 | 0.514425 | 3.94E-05 | 16 | Gas6 | 0.462585 | 0.000359 |
| 7 | Plekha4 | 0.495353 | 9.77E-05 | 17 | Mt3 | 0.459751 | 0.000400 |

| 8 | Nr4a1 | 0.494369 | 9.77E-05 | 18 | Pdlim1 | 0.456149 | 0.000467 |
| --- | --- | --- | --- | --- | --- | --- | --- |
| 9 | Cilp | 0.494264 | 9.77E-05 | 19 | Scn7a | 0.452321 | 0.000552 |
| 10 | Selenop | 0.490979 | 0.000109 | 20 | Mal | 0.442638 | 0.000866 |
| **g:Profiler**  **Extracellular space:** Ctsk, Lbp, Sparcl1, Apod, Clip, Selenop, Aqp1, Igfbps, Gas6  p_val_adj = 4.985E-03 | | | | | | | |

**Supplementary Table 3. List of top 20 DEGs sorted by FC in cluster 1 in subgroup analysis.**

| **Order** | **Gene** | **log2FC** | **p_val_adj** | **Order** | **Gene** | **log2FC** | **p_val_adj** |
| --- | --- | --- | --- | --- | --- | --- | --- |
| 1 | Hbb-bs | 1.481838 | 3.72E-11 | 11 | mt-Nd2 | 1.174465 | 3.11E-18 |
| 2 | Hba-a1 | 1.371583 | 4.03E-08 | 12 | mt-Nd3 | 1.162475 | 1.92E-15 |
| 3 | Hba-a2 | 1.337731 | 6.26E-08 | 13 | Col3a1 | 1.159569 | 2.32E-16 |
| 4 | mt-Nd1 | 1.328159 | 6.40E-26 | 14 | mt-Co2 | 1.158804 | 1.56E-23 |
| 5 | Fabp4 | 1.286137 | 1.71E-09 | 15 | Gpx3 | 1.063088 | 2.04E-12 |
| 6 | Mpz | 1.283254 | 0.002548 | 16 | Apoe | 1.057814 | 7.24E-16 |
| 7 | Apod | 1.245959 | 4.29E-07 | 17 | Mt-Atp6 | 1.004673 | 2.14E-25 |
| 8 | mt-Nd4 | 1.20987 | 2.21E-22 | 18 | Mt-Co3 | 1.000775 | 8.79E-24 |
| 9 | Aqp1 | 1.185255 | 3.18E-06 | 19 | Sparcl1 | 0.996717 | 6.15E-05 |
| 10 | mt-Cytb | 1.179101 | 2.18E-26 | 20 | Plvap | 0.953745 | 0.003773 |
| **g:Profiler**  **Respiratory electron transport chain:** Mt-Nd1, Mt-Nd4, Mt-Cytb, Mt-Nd2, Mt-Nd3, Mt-Co2, Mt-Co3  p_val_adj = 1.885E-09  **Hemoglobin complex:** Hbb-bs, Hba-a1, Hba-a2  p_val_adj = 1.478E-05  **Extracellular space:** Hbb-bs, Hba-a1, Hba-a2, Mt-Nd1, Apod, Aqp1, Col3a1, Gpx3, Apoe, Sparcl1  p_val_adj = 4.985E-03 | | | | | | | |

**Supplementary Table 4. List of top 20 DEGs sorted by FC in cluster 2 in subgroup analysis.**

| **Order** | **Gene** | **log2FC** | **p_val_adj** | **Order** | **Gene** | **log2FC** | **p_val_adj** |
| --- | --- | --- | --- | --- | --- | --- | --- |
| 1 | Ndrg1 | 0.890011 | 5.91E-10 | 11 | Mmp12 | 0.511167 | 0.000615 |
| 2 | Espn | 0.669112 | 3.85E-13 | 12 | Higd1a | 0.506683 | 1.02E-06 |
| 3 | Ero1l | 0.647370 | 3.45E-07 | 13 | Plod1 | 0.475063 | 8.55E-06 |
| 4 | Car9 | 0.594415 | 5.85E-10 | 14 | Pfkp | 0.462048 | 9.69E-08 |
| 5 | Bnip3 | 0.581541 | 3.93E-07 | 15 | Vegfa | 0.459540 | 1.48E-05 |
| 6 | Slc2a1 | 0.553518 | 1.11E-05 | 16 | 2410006H16Rik | 0.452441 | 5.50E-06 |
| 7 | Lgals3 | 0.548944 | 5.84E-08 | 17 | Fam129b | 0.446521 | 0.000179 |
| 8 | Egln1 | 0.545318 | 1.87E-06 | 18 | Bnip3l | 0.442225 | 0.000341 |
| 9 | Gapdh | 0.525084 | 8.00E-15 | 19 | Mif | 0.442142 | 4.57E-13 |
| 10 | Plod2 | 0.514761 | 4.97E-07 | 20 | Ftl1 | 0.437711 | 1.27E-08 |
| **g:Profiler**  **Response to hypoxia:** Ndrg1, Car9, Bnip3, Slc2a1, Egln1, Plod2, Hidg1a, Plod1, Vegfa, Bnip3l  p_val_adj = 1.741E-11  **Glycolysis and gluconeogenesis:** Slc2a1, Gapdh, Pfkp  p_val_adj = 1.196E-03  **Negative regulation of apoptotic process:** Bnip3, Lgals3, Gapdh, Higd1a, Vegfa, Bnip3l, Mif  p_val_adj = 8.615E-03 | | | | | | | |

**Supplementary Table 5. DEGs with significant correlation in cluster 2.**

| **Order** | **Gene** | **Correlation Coefficient** | **P value** | **Order** | **Gene** | **Correlation Coefficient** | **P value** |
| --- | --- | --- | --- | --- | --- | --- | --- |
| 1 | Vdac1 | 0.338128 | 0.000477 | 17 | Higd1a | 0.227459 | 0.020852 |
| 2 | Bnip3 | 0.323650 | 0.000854 | 18 | Plod2 | 0.224801 | 0.022434 |
| 3 | Pgm1 | 0.321714 | 0.000921 | 19 | Aldoa | 0.222327 | 0.023998 |
| 4 | Ero1l | 0.300824 | 0.002018 | 20 | Espn | 0.217574 | 0.027265 |
| 5 | Pfkp | 0.291526 | 0.002810 | 21 | Gys1 | 0.216986 | 0.027694 |
| 6 | Ugdh | 0.288018 | 0.003176 | 22 | Bsg | 0.213404 | 0.030436 |
| 7 | Selenbp1 | 0.281629 | 0.003953 | 23 | Hmga2 | 0.213075 | 0.030698 |
| 8 | Car9 | 0.261373 | 0.007658 | 24 | Cda | 0.209064 | 0.034061 |
| 9 | Gbe1 | 0.257557 | 0.008627 | 25 | Pgk1 | 0.209030 | 0.034091 |
| 10 | Rara | 0.253615 | 0.009740 | 26 | Tnfrsf9 | 0.206881 | 0.036018 |
| 11 | Tpi1 | 0.238939 | 0.015070 | 27 | Ndufv3 | 0.205383 | 0.037415 |
| 12 | Fam162a | 0.237865 | 0.015545 | 28 | Fam129b | 0.200801 | 0.041972 |
| 13 | Ier3 | 0.233566 | 0.017575 | 29 | Ankrd37 | 0.200272 | 0.042527 |
| 14 | Krt19 | 0.232929 | 0.017895 | 30 | Vegfa | 0.198347 | 0.044597 |
| 15 | Ndrg1 | 0.231629 | 0.018563 | 31 | Smox | 0.195299 | 0.048046 |
| 16 | Lgals3 | 0.228386 | 0.020323 |  |  |  |  |
| **g:Profiler**  **Response to hypoxia:** Bnip3, Car9, Fam162a, Ndrg1, Higd1a, Plod2, Aldoa, Pgk1, Vegfa  p_val_adj = 6.079E-07  **Glycolysis and gluconeogenesis:** Pgm1, Pfkp, Tpi1, Aldoa, Pgk1  p_val_adj = 2.381E-05  **Negative regulation of apoptotic process:** Vdac1, Bnip3, Gbe1, Rara, Ier3, Lgals3, Higd1a, Hmga2, Vegfa  p_val_adj = 7.423E-03 | | | | | | | |
